# Using reciprocity for relating the simulation of transcranial current stimulation to the EEG forward problem

**DOI:** 10.1101/043554

**Authors:** S. Wagner, F. Lucka, J. Vorwerk, C.S. Herrmann, G. Nolte, M. Burger, C.H. Wolters

## Abstract

To explore the relationship between transcranial current stimulation (tCS) and the electroencephalography (EEG) forward problem, we investigate and compare accuracy and efficiency of a reciprocal and a direct EEG forward approach for dipolar primary current sources both based on the finite element method (FEM), namely the adjoint approach (AA) and the partial integration approach in conjunction with a transfer matrix concept (PI). By analyzing numerical results, comparing to analytically derived EEG forward potentials and estimating computational complexity in spherical shell models, AA turns out to be essentially identical to PI. It is then proven that AA and PI are also algebraically identical even for general head models. This relation offers a direct link between the EEG forward problem and tCS. We then demonstrate how the quasi-analytical EEG forward solutions in sphere models can be used to validate the numerical accuracies of FEM-based tCS simulation approaches. These approaches differ with respect to the ease with which they can be employed for realistic head modeling based on MRI-derived segmentations. We show that while the accuracy of the most easy to realize approach based on regular hexahedral elements is already quite high, it can be significantly improved if a geometry-adaptation of the elements is employed in conjunction with an isoparametric FEM approach. While the latter approach does not involve any additional difficulties for the user, it reaches the high accuracies of surface-segmentation based tetrahedral FEM, which is considerably more difficult to implement and topologically less flexible in practice. Finally, in a highly realistic head volume conductor model and when compared to the regular alternative, the geometry-adapted hexahedral FEM is shown to result in significant changes in tCS current flow orientation and magnitude up to 45 degrees and a factor of 1.66, respectively.

## 1 Introduction

Brain stimulation techniques such as transcranial current stimulation (tCS) have gained significant impact in the treatment of neuropsychiatric diseases such as Alzheimer’s disease [1], Parkinson’s disease [2] and epilepsy [3]. For tCS, a weak direct current (0.25-2 mA) is injected via one or more electrodes (anode) attached to the scalp and extracted from at least another one (cathode). Depending on the polarity of the currents, neural activity can be modulated. It is important to investigate the underlying mechanisms of tCS in experimental studies, however, such studies are difficult to parametrize, time-consuming and expensive, e.g., because of the possible need for individualized electrode setups and injected current patterns [4].

Computer simulation studies are relatively cheap to perform and allow a deep analysis of the individual current flow in the brain. Consequently, current flow patterns have been predicted in spherical shell models [5, 6] and MRI-derived models of healthy subjects [7, 4, 8, 9] and patients [10, 11, 12]. With regard to head volume conductor modeling, in [9] we investigated the influence of the most important tissue compartments to be modeled in tCS computer simulation studies and presented a guideline for efficient yet accurate volume conductor modeling. We recommended to accurately model the major tissues between the stimulating electrodes and the target areas, while for efficient yet accurate modeling, an exact representation of other tissues seemed less important. We found that at least appropriate isotropic representations of the compartments skin, skull, CSF, gray and white matter seemed to be indispensible. We recommended that, when a significant part of spongy bone is in between the stimulating electrodes and the target region, the distinction between skull compacta and spongiosa should also be added to a realistic head model. Furthermore, white matter anisotropy modeling seemed important only for deeper target regions. Even though the predicted current flow patterns often fit to experimental results [8] and even allow to calculate individually optimized stimulation protocols [4, 13], the reliability of computer simulation studies needs to be validated by its own. One important source of errors, which has not yet sufficiently been investigated in tCS modeling, are numerical errors that may strongly influence the simulation results. In this paper, we will study numerical errors of finite element (FE) approaches for tCS modeling.

In the literature, tetrahedral [5, 14] as well as regular hexahedral [10, 11, 7] FE approaches have been used for tCS modeling. Constrained Delaunay tetrahedralizations (CDT) from given tissue surface representations [15, 16] have the advantage that smooth tissue surfaces are well represented in the model, while on the other side, the generation of such models is difficult in practice and might lead to unrealistic model features. For example, holes in tissue compartments such as the foramen magnum and the optic canals in the skull are often artificially closed to allow CDT meshing. Furthermore, CDT modeling necessitates nested surfaces, while, in reality, surfaces might touch each other like for example the inner and outer surface of the cerebrospinal fluid (CSF) compartment. Hexahedral models do not suffer from such limitations and can be easily generated from voxel-based magnetic resonance imaging (MRI) data, but smooth tissue boundaries cannot be well represented in regular hexahedral models. Therefore, we recently presented an isoparametric FE approach using geometry-adapted hexahedra for tCS simulations [9]. However, to the best of our knowledge, there is no study yet investigating the effect strength of numerical errors of such FE approaches for tCS. Therefore, it is one goal of this study to numerically validate the different FE approaches in a simplified geometry.

In contrast, in the related field of EEG source analysis, validation of numerical EEG forward modeling approaches has been carried out in simplified geometries such as multilayer-sphere models, where quasi-analytical series expansion formulas for the EEG forward problem have been derived (see recent review in [15]).

In this work, two approaches for the EEG forward problem are employed. While the partial integration approach [17] uses integration-by-parts to solve the EEG forward problem in a direct manner, the adjoint approach [18] applies the adjoint method to compute the EEG forward potential from a sensor-point of view. In both approaches, the source terms are assumed to be square-integrable functions on the head domain. In source analysis, however, most often an equivalent current dipole is used as a primary current source [15], which does not fulfill these requirements so that the regularity assumptions in the derivations of PI and AA are not fulfilled (see deeper discussion of this issue in the theory section 2.1). It is thus also one aim of this study to investigate in sphere models whether the use of an equivalent current dipole in the adjoint partial differential equation leads to different numerical errors when compared to the direct partial integration approach.

Our overall validation strategy for FE-based tCS modeling in this paper is to construct a theoretical link between tCS and EEG forward modeling, so that numerical results in simplified volume conductor models can be exploited for validation of both applications. Therefore, we will first introduce the EEG forward problem (Section 2.1) and, for its solution, the partial integration approach (Section 2.2) in conjunction with the FE transfer matrix concept (Section 2.3), whose combination will be abbreviated in the following by just PI. The adjoint approach (AA, Section 2.4) is then presented as a method that offers a close link between the EEG forward problem and tCS modeling (Section 2.5). The remainder of Section 3 will report on the different tetrahedral and hexahedral FE approaches used in this study, the validation platform for the spherical model validations (Section 3.1) and an overview on how we built the volume conductor for the realistic head model evaluation study (Section 3.2). Section 4 presents the results of our study. We first show in CDT and in regular and geometry-adapted hexahedral multilayer-sphere FE models that AA and PI for dipolar primary current sources lead to identical numerical errors in comparison to quasi-analytical series expansion formulas and to a nearly identical computational complexity (Section 4.1). In the Appendix A, we prove that AA and PI for dipolar primary current sources are algebraically identical even for general head models. We thus derive a theoretical relationship between the EEG forward problem and tCS simulation. Finally, using a realistic volume conductor model and a commonly used electrode arrangement for auditory cortex stimulation, we investigate current flow vector field orientation and magnitude changes between a regular and a geometry-adapted hexahedral FE approach for tCS simulation (Section 4.2). In 5, we discuss the results, summarize the most important findings and conclude our study.

## 2 Theory

### 2.1. EEG forward problem

Applying the quasi-static approximation of Maxwell’s equations [19] for computing the electric potential Φ for a primary current density in the brain, ***J***^*p*^, yields a Poisson equation with homogeneous Neumann boundary conditions [15]

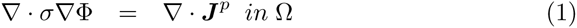

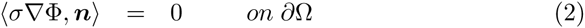

with *σ* : ℝ^3^ → ℝ^3×3^ being a 3 × 3 tensor of tissue conductivity, *n* being the outer unit normal at the scalp surface and Ω and ∂Ω being the head domain and its boundary, respectively.

The primary current density ***J***^*p*^ is commonly modeled as a mathematical point-dipole source at location *x*_0_ with moment *q* [20, 15]:

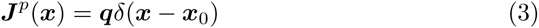

The source term in both following approaches, i.e., in section 2.2 the partial integration approach [17] and in section 2.4 the adjoint approach [18], is assumed to be a square-integrable function on the head domain Ω, i.e., it is assumed to be in the Sobolev space *H*^0^(Ω) = *L*^2^(Ω). However, later in the derivations, a current dipole is then assumed as source term. As shown in equation (3), a dipole is made of a Dirac distribution *δ* in 3-dimensional space, which is only in the Sobolev space *H*^‒3/2‒*ε*^(Ω) (and thus not in *L*^*2*^(Ω)) [21, 16], so that the right-hand side of the PDE in equation (1), i.e., the divergence of a dipole, loses even one more degree of regularity and is thus only in the Sobolev space *H*^‒5/2–ε^(Ω) [21]. Therefore, the regularity assumptions with regard to the source term and the resulting right hand sides of the PDEs in the derivations of both approaches are not fulfilled. The numerical comparison of the two approaches in the presence of dipolar primary current sources is thus of great interest.

### 2.2. Partial integration approach

Here, we summarize the most important steps for the derivation of the partial integration FE approach. A more detailed derivation can be found in [17, 22, 15]. Firstly, the partial differential equation (PDE) for the EEG forward problem (1) is multiplied with a FE ansatz function *ϕ*_*i*_ and integrated over Ω. Secondly, integration by parts is applied on both sides. Thirdly, boundary condition (2) is exploited and finally the potential Φ is projected into the FE space. This leads to the linear equation system

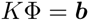

with the high-dimensional, but sparsely populated *stiffness matrix K*

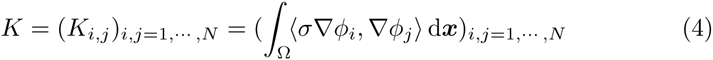

and the right-hand side vector *b*

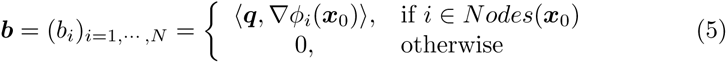

with *N* being the number of FE nodes and *Nodes*(*x*_0_) defining the nodes of the elements containing the dipole position.

### 2.3. FE transfer matrix approach

The transfer matrix approach for FE-based EEG source analysis was introduced in [23] and extended to magnetoencephalography (MEG) in [24].

When solving the PDE (1,2) for an arbitrary dipolar source, the potential Φ is calculated in the whole volume conductor. However, with regard to the EEG inverse problem, one is only interested in the potential at the few FE nodes that are identified with the electrode positions. Thus, for the partial integration approach, we introduce a restriction matrix *R* ∈ ℝ^(*S*–1)×*N*^, mapping Φ to the *S* – 1 non reference electrodes

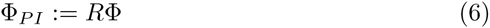

We can then define the *transfer matrix T* as follows:

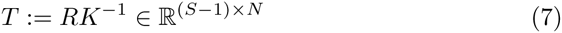

The calculation of *T* requires to solve only *S* – 1 large, but sparse FE equation systems, which can be done very efficiently using an algebraic multigrid preconditioned conjugate gradient (AMG-CG) solver that is specifically tailored for our problem in inhomogeneous and anisotropic head volume conductor models [25, 26]. Furthermore, in many practical applications *T* can be pre-computed, enabling a fast and robust computation of many forward solutions whenever desired. Once the transfer matrix is calculated, the solution Φ_*EEG*_ of the EEG forward problem is given by the product of *T* with the sparse right-hand side vector *b* from equation (5):

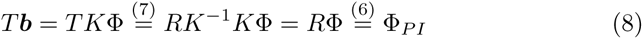

In the following, we will call the combination of the partial integration method from Section 2.2 with the above FE transfer matrix approach the PI approach for the EEG forward problem.

### 2.4. Adjoint approach

The reciprocal or adjoint approach [27, 28, 29, 30, 18] switches the role of the sources with the sensors, resulting in a Laplace equation with inhomogeneous Neumann boundary conditions

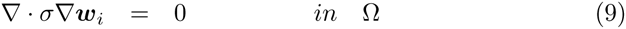

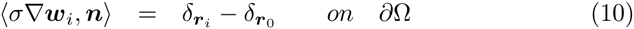

with *r*_*i*_ and *r*_0_ being the positions of the *i*-th surface electrode and the reference electrode, respectively. For numerical realization of the adjoint approach, Equation (9) is multiplied with a FE ansatz function *ϕ*_*j*_ and integrated over Ω. Next, integration by parts is applied to the left-hand side and the boundary condition (10) is used. Finally, *w*_*i*_ is projected into the FE space, leading to the linear equation system

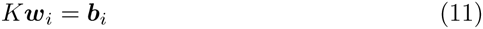

with the Stiffness matrix (4) and the right-hand side vector

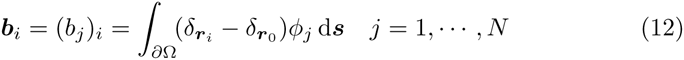

After solving this equation, a combination of Riesz representation theorem and Helmholtz’ principle of reciprocity relates the solution *w*_*i*_ of the adjoint approach for a current dipole at location *x*_0_ with dipolar moment *q* to the EEG forward potential difference between electrode *i* and the reference electrode [29, 18]:

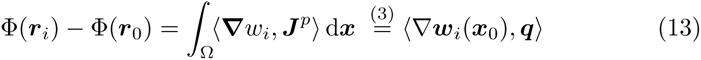

In order to calculate the adjoint approach solution Φ_*AA*_ of the EEG forward problem, the linear equation system (11) has to be solved for *i* = 1, …, *S* – 1 different right-hand sides (12) using for example an AMG-CG, i.e., in each step, we set *b*(*r*_*i*_) = 1 and *b*(*r*_0_) = – 1 and use Equation (13) to calculate the potential differences Φ(*r*_*i*_) – Φ(*r*_0_). Finally, a common average reference is used to obtain the full vector of EEG forward potentials, i.e., we use the additional constraint 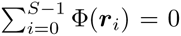. Therefore, the EEG forward potential at the reference electrode can be calculated as 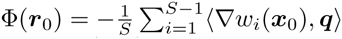. In the following, we will call this the adjoint approach (AA) for the EEG forward problem.

### 2.5. Relationship of the adjoint approach to transcranial current stimulation

For a given pair of reference electrode at *r*_0_ and stimulating electrode at *r*_*i*_, the AA also allows to calculate a so-called *electrode lead vector field* ***S***_*i*_ : ℝ^3^ → ℝ^3^ [27, 29, 18] as

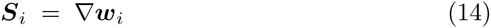

which can be used to visualize the sensitivity of this electrode pair to sources in the brain in just a single image for ***S***_*i*_ and having Equation (13) in mind [31]. More importantly, for relating the AA to tCS simulation, the solution vector *w*_*i*_ of the adjoint PDE (9) and (10) can additionally be used to simulate a current density vector field ***J*** : ℝ^3^ → ℝ^3^ for a current injected at a point electrode *r*_*i*_ and removed at another point electrode *r*_0_ as:

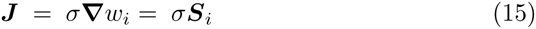

In tCS simulations, however, point-like sensors are not used commonly, but the size and shape of the electrodes and the total current applied to the electrodes (here 1 mA) are taken into account. Therefore, the boundary condition (10) is exchanged by inhomogeneous Neumann boundary conditions at the electrode surfaces and homogeneous Neumann boundary conditions at the remaining model surface. The *tCS forward problem* is thus given as

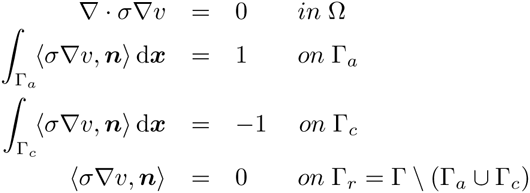

with Γ_*a*_ and Γ_*c*_ being the surfaces of anode and cathode, respectively. When numerically solving this PDE, we again gain a linear equation system *Kv* = *b* with *K* being the stiffness matrix (4), *v* the vector with the potential values at the FE nodes and *b* the right-hand side vector with non-zero entries only at the electrode nodes. This equation system can again be efficiently solved using for example AMG-CG. Finally, the current density ***J*** is computed by multiplying the gradient of the potential field with the conductivity tensor, following (15).

## 3 Methods

### 3.1. Validation platform

#### 3.1.1. Four compartment sphere model

Numerical validation will be performed in a four compartment (scalp, skull, CSF and brain) sphere model using radii and conductivities as indicated in Table 1. Conductivity values are identical to those used in [32, 33]. *S* = 748 EEG electrodes were equally distributed over the surface of the outer sphere. 77 tangential and 77 radial dipoles were placed on a ray from the center of the sphere to the brain/CSF interface in steps of 1 mm. Thus, the most eccentric dipoles are only 1 mm away from the first conductivity jump. For each dipole, the source eccentricity is defined as the percentage of the difference between the dipole location and the sphere midpoint divided by the radius of the innermost shell. The most eccentric dipole thus has an eccentricity of 98.72%.

**Table 1:** Parameterization of the four-layer sphere model.

| Tissue | Scalp | Skull | CSF | Brain |
| --- | --- | --- | --- | --- |
| Outer shell radius (mm) | 92 | 86 | 80 | 78 |
| Conductivity (S/m) | 0.43 | 0.0042 | 1.79 | 0.33 |

#### 3.1.2. FEM mesh generation

In the following, we will shortly summarize the main aspects and parameterizations of the three different FE approaches used in this study, namely constrained Delaunay tetrahedralization, regular and geometry-adapted hexahedral FE approaches.

*Constrained Delaunay tetrahedralization FE approach:* Tetrahedral FE meshes of the four layer sphere model are generated using the software TetGen [34] which uses a constrained Delaunay tetrahedralization (CDT) approach [35, 36]. Using models generated with this approach, EEG source analysis was performed [25, 16]. The meshing procedure starts with the preparation of a suitable boundary discretization of the model. For each of the four layers and for a given triangle *edge* length, nodes are distributed in a most-regular way and connected through triangles. For model *tet503k* of this study, we used an edge length of 1.75 mm. This yields a valid triangular surface mesh for each of the four layers. Meshes of different layers are not allowed to intersect each other. The CDT approach is then used to construct a tetrahedralization conforming to the surface meshes. It first builds a Delaunay tetrahedralization of the vertices of the surface meshes. It then uses a local degeneracy removal algorithm combining vertex perturbation and vertex insertion to construct a new set of vertices which include the input set of vertices. In the last step, a fast facet recovery algorithm is used to construct the CDT [35, 36]. This approach is combined with two further constraints to the size and shape of the tetrahedra. The first constraint can be used to restrict the volume of the generated tetrahedra in a certain compartment, the so-called *volume* constraint. For *tet*503*k*, we used a volume of 0.63 mm^3^. The second constraint is important for the quality of the generated tetrahedra. If R denotes the radius of the unique circumsphere of a tetrahedron and L its shortest edge length, the so-called radius-edge ratio of the tetrahedron can be defined as 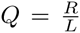. For *tet*503*k*, we used *Q* = 1.0. The radius-edge ratio can detect almost all badly-shaped tetrahedra except one type of tetrahedra, so-called slivers. A sliver is a very flat tetrahedron which has no small edges, but can have arbitrarily large dihedral angles (close to π). For this reason, an additional mesh smoothing and optimization step was used to remove the slivers and improve the overall mesh quality. The CDT meshing procedure for model *tet*503*k* resulted in a tetrahedral mesh with 503 thousand nodes and 3.07 million elements.

*Regular and geometry adapted, hexahedral FE approaches:* While regular hexahedral FE approaches have been used in many simulation studies [10, 11, 7], we developed an isoparametric FE approach for EEG source analysis [37] and for tCS modeling [9] that is specifically tailored to geometry-adapted hexahedral models. First, a hexahedral FE mesh was constructed out of the labeled volume of the four compartment sphere model using an edge length of 1 mm. This resulted in the regular hexahedral FE model *hex*3.2*m* with 3.2 million nodes and 3.24 million elements. To increase conformance to the real geometry and to mitigate the staircase effects of the regular hexahedral mesh, we applied a technique to shift nodes on material interfaces [38, 37]. We chose a nodeshift factor of 0.33, which ensured that the element remained convex and the Jacobian determinant in the FEM computations remained positive in all models (especially also the realistic head model). This procedure resulted in the geometry-adapted hexahedral FE model *gahex3.2m* with 3.2 million nodes and 3.24 million elements. The freely available software SimBio-VGRID^1^ was used for hexahedral mesh generation.

While the hexahedral grid has about 6 times as many nodes as the tetrahedral grid, the number of elements is nearly identical for both approaches for the purpose of similar geometry representation. Since conductivity is modeled constant over each element, the number of elements is a good indicator for the quality of geometry approximation of each model.

#### 3.1.3. FEM modeling

For all FEM computations, we used the freely available SimBio^2^ software toolbox. For the tetrahedral approach, we used linear and for both hexahedral approaches trilinear FE ansatz functions. All equation systems were solved using AMG-CG with solver accuracy of 10^‒7^ (relative error in *KC*^‒1^*K* energy norm with *C* the AMG-preconditioner matrix [25, 26]).

#### 3.1.4. Error measures

For the simplified geometry of a multilayer sphere model, quasi-analytical solutions to compute the potential distribution at the electrodes for a dipole in the brain compartment exist [39]. Such a quasi-analytical solution vector with electrode potentials will be denoted by Φ_*ana*_ ∈ ℝ^*S*^. The analytical solution allows us to calculate topography errors *(relative difference measure* (RDM)) and magnitude errors (*magnification factor* (MAG)) of the numerically-computed potential vector Φ_*num*_ ∈ ℝ^*S*^ as follows [25]:

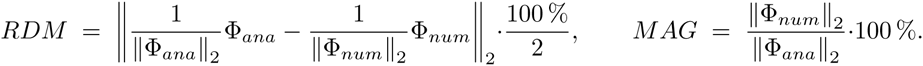

### 3.2. Realistic hexahedral volume conductor models for auditory cortex stimulation

Additionally to the sphere models for validation, we generated highly realistic head volume conductor models for evaluation purposes in order to compare tCS simulation results using the geometry-adapted isoparametric hexahedral FE approach with the simulation results generated in a regular hexahedral model. Therefore, T1-, T2‐ and diffusion-weighted magnetic resonance imaging (MRI) scans of a healthy 26-year-old male subject were measured on a 3T scanner (Gyroscan Intera/Achieva 3.0T, System Release 2.5 (Philips, Best, NL)). A T1w pulse sequence with water selective excitation and a T2w pulse sequence with minimal water-fat-shift were used, both with an isotropic resolution, resulting in cubic voxels of 1.17 mm edge length. This resolution will also define our FE mesh resolution for model *reahead*2, 2*m*. Diffusion weighted MRI was performed using a Stejskal-Tanner spin-echo EPI sequence with a SENSE parallel imaging scheme in the AP direction (acceleration factor 2). Geometry parameters were: FOV 240 mm × 240 mm for 70 transverse slices, 1.875 mm thick, without gap, with a square matrix of 128, resulting in cubic voxels of 1.875 mm edge length. Contrast parameters were TR = 7,546 ms, TE = 67 ms. One volume was acquired with diffusion sensitivity b = 0 s mm^‒2^ and 20 volumes with b = 1,000 s mm^‒2^ using diffusion weighted gradients in 20 directions, equally distributed on a sphere according to the scheme of [40]. The pixel bandwidth was 2,873 Hz/pixel and the bandwidth in the phase-encoding direction was 20.3 Hz/pixel. An additional volume with flat diffusion gradient, i.e., b = 0 s mm^‒2^, was acquired with reversed encoding gradients for later use in susceptibility correction.

The T2w MRI was registered onto the T1w MRI using a rigid registration approach and mutual information [41] as a cost-function as implemented in FSL^3^. Then, the brain, inner skull, outer skull and skin masks were obtained from the T1w and T2w images. In the next step, the T1w image served for the segmentation of gray and white matter and the T2w image for the segmentation of the CSF. For all of these steps, the FSL software was used [42]. The segmentation was visually inspected and manually corrected using CURRY^4^. Segmentation of the skull spongiosa was based on the T2w image. The skull was first constrained using the inner and outer skull masks on the T2w MRI and then a one-voxel-erosion was performed on the skull compartment (this will later guarantee that inner and outer skull compacta are at least one voxel thick [43]). Finally, a thresholding based region-growing segmentation constraint was used to the eroded skull compartment to differentiate between spongiosa and compacta again using CURRY.

Two tCS electrodes were modeled as rectangular patches with a commonly used size of 7 cm × 5 cm [3, 44], thickness of 4 mm, a conductivity value for saline of 1.4 S m^‒1^ [45, 11] and a total current of 1 mA was applied. To simulate the current flow in tCS auditory cortex stimulation, the patches were positioned symmetrically around the auditory cortices, as can be seen in Figure 1.

**Figure 1:**
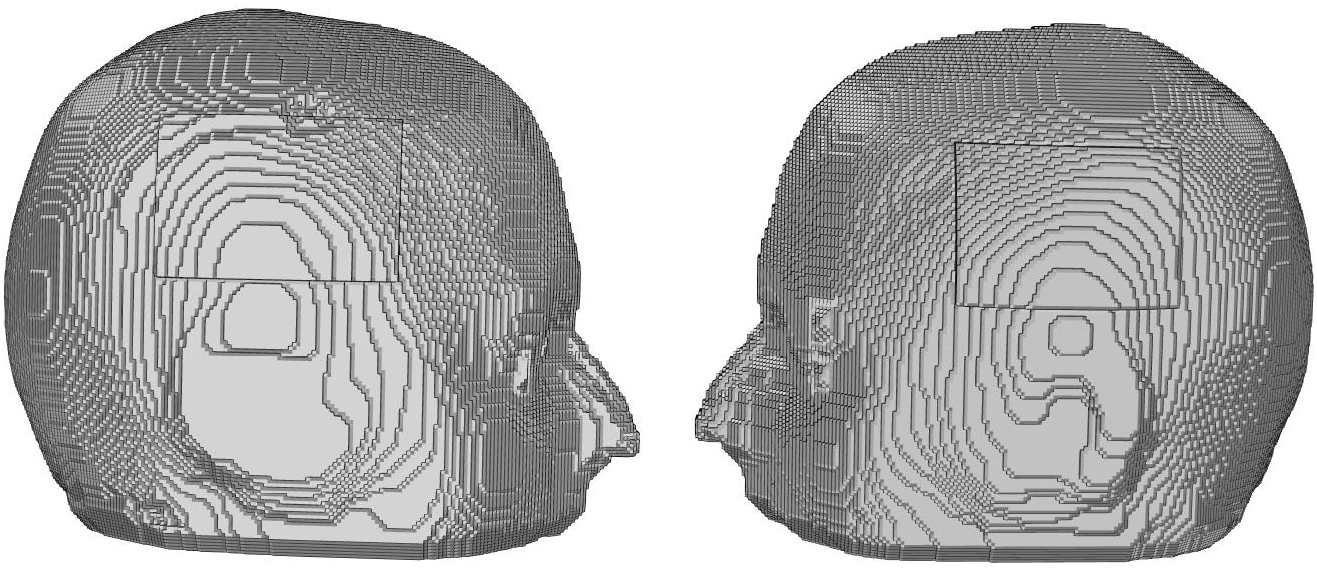
Stimulating electrodes for auditory cortex stimulation. The anode (left subfigure) and cathode (right subfigure) are presented on the model surface.

A regular hexahedral model *head – reghex*2.2*m* and a geometry-adapted hexahedral model *head – gareghex*2.2*m* were generated as described in Section 3.1.2 resulting in meshes with 2.2 million nodes and 2.2 million elements. Tissue conductivity values are identical to those used in [33, 46, 43] and assigned as shown in Table 2.

**Table 2:** Parameterization of the realistic head model.

| Tissue | Scalp | Compacta | Spongiosa | CSF | GM | WM |
| --- | --- | --- | --- | --- | --- | --- |
| Conductivity (S/m) | 0.43 | 0.007 | 0.025 | 1.79 | 0.33 | aniso |

In [47], a model was introduced with which conductivity tensors can be estimated from non-invasively measured DT-MRI. Positive validations of this model were reported by [47] and [48]. We followed this procedure with regard to the white matter compartment as described in the following. We corrected the diffusion-weighted (DW) MR images for eddy current (EC) artifacts by affine registration of the directional images to the b0 image using the FSL routine FLIRT. After this procedure, the gradient directions were reoriented using the rotational part of the transformation matrices obtained in the EC correction scheme. Then, a diffeomorphic approach for the correction of susceptibility artifacts using a reversed gradient approach and multiscale nonlinear image registration was applied to the DW-MRI datasets [49], as implemented in the freely available SPM^5^ and FAIR^6^ software packages. After EC and susceptibility correction, the b0 image was rigidly registered to the T2w image using FLIRT and the transformation matrix obtained in this step was used for the registration of the directional images, while taking care that the corresponding gradient directions were also reoriented accordingly. The tensors were then calculated using the FSL routine DTIFIT [42]. In the last step, white matter conductivity tensors were calculated from the artifact-corrected and registered diffusion tensor MR images using the effective medium approach as described in [47] and embedded in the hexahedral FE head models. The scaling factor between diffusion and conductivity tensors was selected so that the arithmetic mean of the volume of all white matter conductivity tensors optimally fits the volume of the isotropic approximation in a least squares sense [50].

## 4 Results

### 4.1. Validation in four layer sphere model

Figures 2 and 3 show RDM and MAG errors for the partial integration in combination with the FE transfer matrix approach (PI) and the adjoint approach (AA) using CDT model *tet*503*k*, the regular hexahedral model *hex*3.2*m* and the geometry-adapted hexahedral model *gahex*3.2*m*.

**Figure 2:**
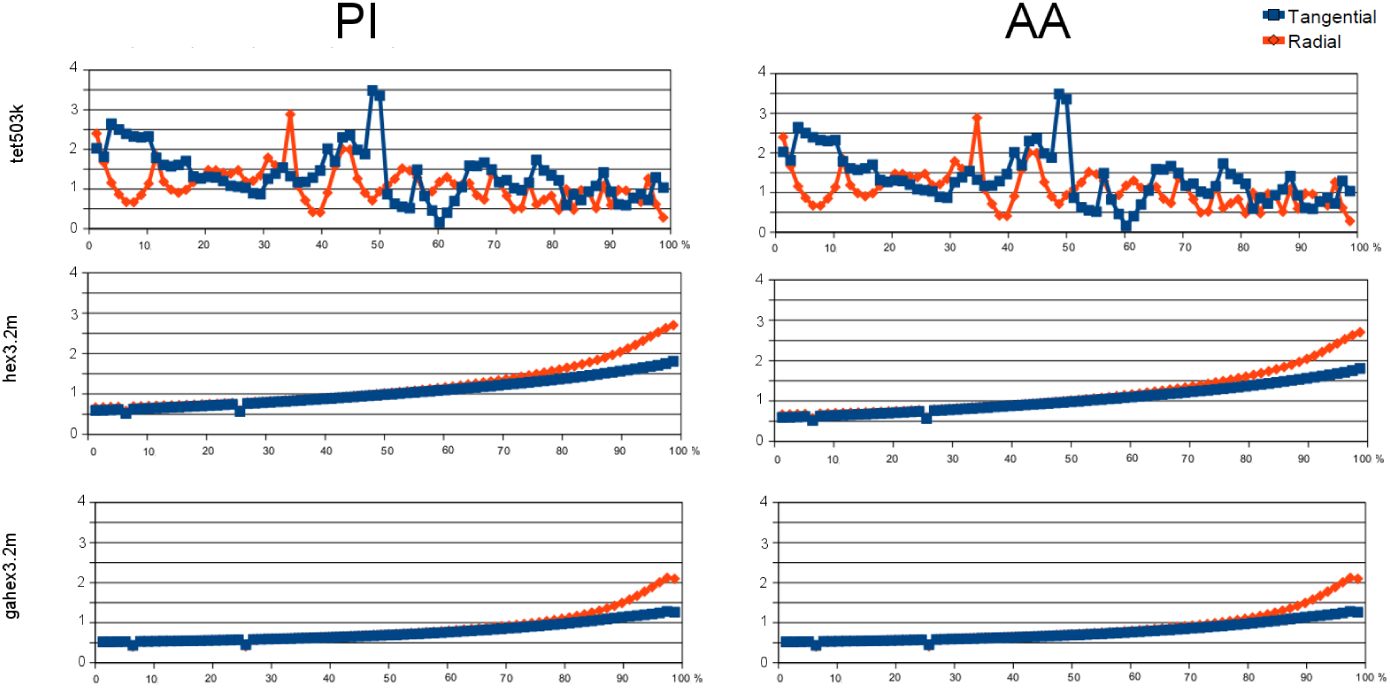
Validations in spherical shell model: RDM errors for PI (left column) and AA (right column) in CDT model *tet*503*k* (first row), in regular hexahedral model *hex*3.2*m* (second row) and in geometry-adapted hexahedral model *gahex*3.2*m* (third row) at different source eccentricities (x-axis). Errors for tangential (blue) and radial sources (red) are color-coded. Note that the scaling on the y-axes is identical to allow an easy comparison of numerical errors.

**Figure 3:**
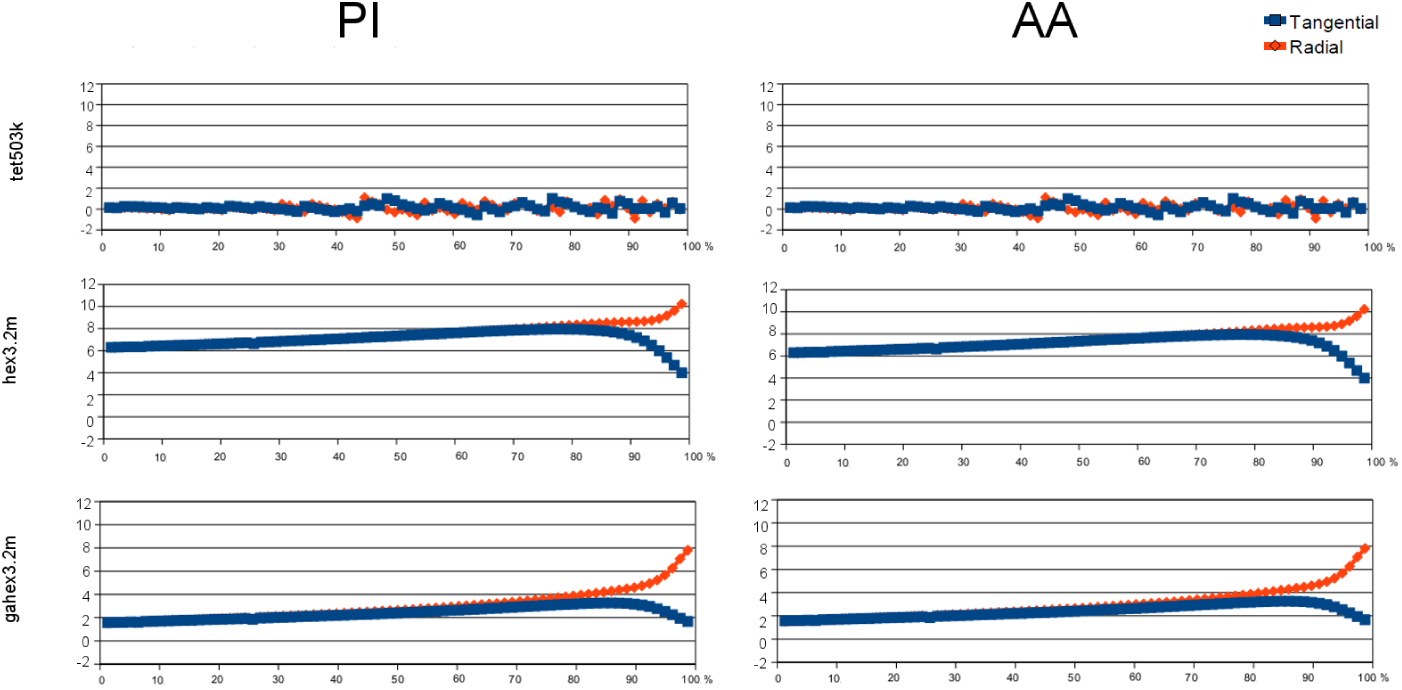
Validations in spherical shell model: MAG errors for PI (left column) and AA (right column) in CDT model *tet*503*k* (first row), in regular hexahedral model *hex*3.2*m* (second row) and in geometry-adapted hexahedral model *gahex*3.2*m* (third row) at different source eccentricities (x-axis). Errors for tangential (blue) and radial sources (red) are color-coded. Note that the scaling on the y-axes is identical to allow an easy comparison of numerical errors.

Most importantly and as can be seen in both figures, AA and PI perform identically with respect to numerical accuracy, independent of depth and orientation of sources and number, size and shape of elements to be considered. The differences in potential solutions are in the range of the solver accuracy (see section 3.1.3). For the interplay between chosen solver accuracy, the discretization error and the numerical accuracy for the PI approach, we refer to [25]. The arithmetic operations count (see also [31]) in Table 3 shows that PI and AA require the same amount of operations to solve the EEG forward problem for all three models. The combination of Figures 2, 3 and Table 3 points out that PI and AA are identical for the four-layer sphere scenario, even if the way of computation is quite different.

**Table 3.**
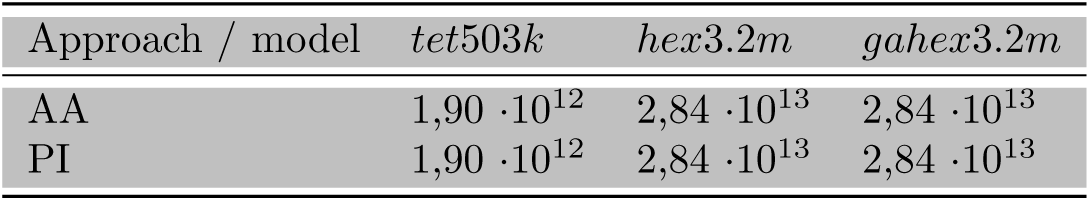
Arithmetic operations needed for PI and AA to solve the EEG forward problem in tetrahedral model *tet*503*k* and in regular and geometry-adapted hexahedral models *hex*3.2*m* and *gahex*3.2*m*, respectively.

In Theorem 1 in the Appendix, we are able to even prove algebraically that, up to the reference potential, PI and AA will lead to the same results, even for general head models. With regard to the reference, PI fixes the potential at a reference electrode to be exactly zero (Φ(r_0_) = 0), while AA calculates the potential at the measurement sites with respect to the potential at a reference electrode (Φ(r_*i*_) – Φ(r_0_)). However, after re-referencing, both PI and AA will lead to the same results (see Appendix).

In Sections 2.4 and 2.5 it was shown that, as a substep of the overall calculations, in equations (11) and (12) the AA numerically computes the electric potential solution for a tCS simulation when using point-electrodes at r**i** and r_0_. Therefore, the validation results presented in Figures 2 and 3 do not only show the numerical errors in the EEG forward problem, but also allow conclusions about the accuracy in calculating the electric potential underlying tCS simulations.

As shown in Figure 2, with regard to the RDM and for eccentricities between 0% and 70%, the geometry-adapted hexahedral approach performs best (≤ 1%) when compared to the regular hexahedral approach (≤ 1.4%) and the CDT approach (≤ 3.5%). For the higher eccentricities between 70% and 98.72%, with RDM errors below 1.8%, the CDT is slightly better than the geometry-adapted (≤ 2.1%) and the regular hexahedral approach (≤ 2.8%). For both hexahedral modeling approaches and for higher eccentricities between 70% and 98.72%, RDM errors are higher for radial than for tangential sources, while they are very similar for eccentricities below 70%.

As shown in Figure 3, with errors below 1.1%, the CDT approach performs best with regard to the MAG, as expected. It is followed by the geometry-adapted hexahedral approach (≤ 4% for eccentricities below 80% and ≤ 8% for excentricities below 98.72%) and the regular hexahedral approach (≤ 8.2% for eccentricities below 80% and ≤ 11% for eccentricities below 98.72%). For both hexahedral modeling approaches and for higher eccentricities between 70% and 98.72%, MAG errors are higher for radial than for tangential sources, while they are very similar for eccentricities below 70%.

As deeper discussed in Section 5, a comparison between the CDT and the hexahedral approaches is rather difficult, while with regard to realistic head models both hexahedral models can directly be compared. Therefore, and as a preparation for Section 4.2, we now summarize our hexahedral model validation results. The geometry-adapted approach outperforms the standard regular approach by a factor of 1.3 with regard to RDM and MAG (from a maximal RDM of 2.8% down to 2.1% and from a maximal MAG of 11% down to 8%), further motivating the use of geometry-adaptation in combination with an isoparametric hexahedral FE approach for practical applications.

### 4.2. Regular and geometry-adapted hexahedral approaches in a realistic tCS model

Figure 4 depicts changes in orientation (top row) and magnitude (bottom row) of the current density vector field (see Equation (15)) in the whole volume conductor (left column) and only in the brain compartment (right column) when using the geometry-adapted hexahedral approach as compared to the standard regular one.

**Figure 4:**
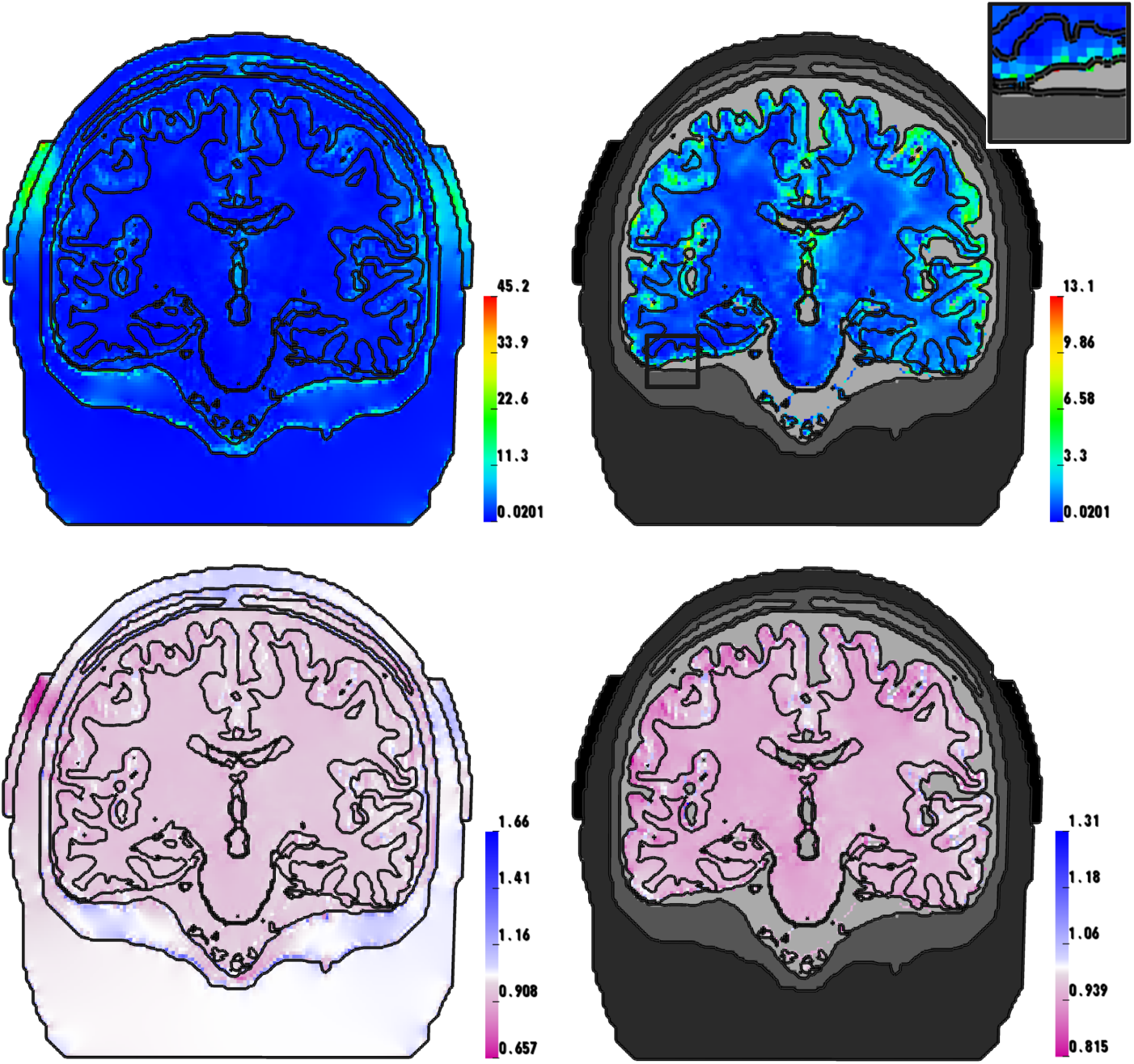
Differences in the resulting current density vector field between an isoparametric geometry-adapted hexahedral FE approach as compared to a standard regular hexahedral approach for auditory cortex stimulation: Changes in orientation (top row) and magnitude (bottom row) in the whole volume conductor (left column) and only in the brain compartment (right column). The subfigure top right shows a magnification of the black box area where the maximum in orientation change is achieved.

As can be seen, with orientation changes up to 45.2 degrees, changes are largest in the electrodes (upper left subfigure). Moreover, significant changes in orientation can be seen in skin, skull and CSF compartments. In superficial cortical areas, we still find maximal current density orientation changes up to 13.1 degrees (upper right subfigure, see especially the magnified black box where the maximum in orientation change is achieved).

The change in magnitude is depicted on a pink-white-blue scale (bottom row), i.e., magnitude in the geometry adapted model as compared to the regular model decreases in pink areas, remains constant in white regions and increases in blue regions. For auditory cortex stimulation, both an increase (up to 66%) and a decrease (up to 44%) in current density magnitude can be seen in electrodes, skin and skull (lower left). Moreover, magnitude mainly decreases in the CSF (lower left) and, with values up to 18.5%, in the brain compartment (lower right).

## 5 Discussion and conclusion

In this study, we investigated accuracy and efficiency of a reciprocal and a direct EEG forward approach, namely the adjoint approach (AA) [27, 28, 29, 30, 18] and the partial integration approach in conjunction with an FE transfer matrix concept (PI) [17, 23, 22, 24]. Since the regularity assumptions in the derivations of both approaches are not fulfilled for a dipolar primary current source, we analyzed the numerical accuracies of both approaches with regard to topography (RDM) and magnitude (MAG) numerical errors in a four compartment sphere model, where quasi-analytical series expansion formulas exist [39]. We used and compared a constrained Delaunay tetrahedralization (CDT) FE approach, a standard hexahedral FE approach based on regular hexahedra and an isoparametric FE approach using geometry-adapted hexahedra.

Our validation study in a multilayer sphere model revealed that the numerical accuracy and computational complexity for calculating the EEG forward problem is nearly identical for the AA and the PI approach for dipolar current sources and that differences are only in the solver accuracy range. Moreover, we could prove algebraically that AA and PI are identical for dipolar sources even for general head models. With regard to improving the numerical accuracy and efficiency of FE-based EEG forward modeling, it is thus sufficient to only validate one of the approaches and compare it with other source modeling approaches [51, 25, 37, 22]. Even if AA thus does not contribute new numerical aspects to the EEG forward problem, it allows to calculate electrode lead vector fields ***S*** for given electrode pairs (see equation (14)) and visualize the sensitivity of this electrode pair to sources in the brain using just a single image for ***S*** and having equation (13) in mind.

Furthermore, as shown in this paper, the AA can also be used to bridge EEG source analysis and tCS simulation (see equations (13), (14) and (15)). Therefore, quasi-analytical EEG forward solutions in multilayer sphere models [39] can not only be used to investigate numerical accuracies of different FE approaches for the EEG forward problem, but also to reciprocally validate the approaches for tCS simulation. For tCS modeling, tetrahedral approaches have been presented in [5, 14], regular hexahedral approaches in [10, 11, 7] and the geometry-adapted hexahedral approach in [9].

As shown in Figures 2 and 3, the geometry-adapted approach is more accurate by a factor of 1.3 than the standard regular hexahedral approach. For the MAG error (Figure 3), an offset between the regular and the geometry-adapted hexahedral approach can be seen. This is due to the fact that the geometry-adapted hexahedral model in combination with an isoparametric FE approach better approximates the real shape of the sphere model, leading to significantly lower magnitude errors. The gain in accuracy by means of a better approximation of smooth compartment surfaces thus outweighs the possible numerical disadvantage of less regular elements at tissue boundaries. The result presented here is in line with the EEG forward simulation study of [37], which showed that topography and magnitude errors can be reduced by even more than a factor of 2 and 1.5 for tangential and radial sources, respectively, when using an isoparametric FE approach with geometry-adapted hexahedral meshes with a nodeshift factor of 0. 49 and Venant and subtraction source modeling approaches in 2 and 3 mm hexahedral models. In our study, besides the higher resolution (1 mm) and the different source modeling approach (PI), we only used a nodeshift factor of 0.33, which explains the smaller factor of error-reduction. The more conservative nodeshift-factor ensures that interior angles at element vertices remain convex and the Jacobian determinants in the FEM computations remain positive also in highly realistic six-compartment head models [9, 52]. We therefore also used it here for the spherical model validation study. In summary, the geometry-adapted approach does not involve any additional difficulties for the user and thus has to be preferred to the regular standard approach for modeling both the EEG forward problem and tCS.

The comparison between the CDT approach and the geometry-adapted hexahedral approach is more difficult. For the purpose of similar geometry representation, we chose a similar number of elements for the tetrahedral (3.07m) and hexahedral (3.24m) modeling approaches. However, first of all, model *gahex*3.2*m* has about 6 times more degrees of freedom when compared to model *tet*503*k*. When knowing that the computational complexity for FE modeling in both source analysis and tCS simulation using an algebraic multigrid conjugate gradient solver (AMG-CG) increases mainly linearly with the number of unknowns, *gahex3.2m* leads to 6 times higher computational costs than tet503k. Even if for eccentricities between 0% and 70%, with ≤ 1% compared to ≤ 3.5% RDM error, the *gahex*3.2*m* based approach outperformed the tet503k based approach, for the higher eccentricities between 70% and 98.72% the less-computationally expensive tetrahedral approach even slightly took the lead with RDM errors ≤ 1.8% compared to ≤ 2.1%. With regard to the MAG, the CDT (≤ 1.1%) outperformed the geometry-adapted hexahedral approach (≤ 4% for eccentricities below 80% and ≤ 8% for excentricities below 98.72%). With regard to multilayer sphere modeling, where non-intersecting surfaces can accurately be constructed, CDT FE approaches are thus advisable. However, with regard to the modeling of a realistic head volume conductor from voxel-based magnetic resonance imaging (MRI) data, numerical errors of a few percent are most probably negligible when compared to remaining model errors. This has been shown by [51], who concluded that a reduction of the model error will have a much higher impact than a further increase of the numerical accuracy. A CDT FE approach for realistic head geometries is difficult to generate in practice and might lead to unrealistic model features like artificially closed skull compartments that ignore skull holes like the foramen magnum and the optic canals. Furthermore, CDT modeling necessitates nested surfaces, while in reality surfaces might touch each other, e.g., the inner and outer surface of the cerebrospinal fluid compartment. As discussed below, it might thus be advisable to focus on further reducing model errors. Therefore, for realistic head modeling, we consider voxel-based methods like the isoparametric geometry-adapted hexahedral FE approach as more advisable than purely surface-based ones because of their convenient generation from MRI data and the accompanying topological advantages. Finally, we would like to mention two interesting future directions that both have the goal to combine the advantages of smooth surfaces given, e.g., by levelset segmentations and the topological advantages of voxel-segmentations, namely the immersed FEM [53] and the unfitted discontinuous Galerkin approach [54].

In Equation (15), we showed that lead vector fields in EEG source analysis and tCS current flow fields are directly related as the latter is identical to the product of the conductivity tensor *σ* and the lead vector field ***S*** (when considering identical electrode montages). Therefore, our results also link head volume conductor model sensitivity investigations in EEG source analysis to tCS modeling and vice versa. For example, in Wagner and colleagues [9], tCS computer simulations were performed, starting with a homogenized three-compartment head model and extending this step by step to a six-compartment anisotropic model. Thereby, important tCS volume conduction effects were shown and a guideline for efficient yet accurate volume conductor modeling was presented. Vorwerk and colleagues [55] investigated the influence of modeling/not modeling the compartments skull spongiosa, skull compacta, CSF, gray matter, and white matter and of the inclusion of white matter anisotropy on the EEG forward solutions. The effect sizes in terms of orientation and magnitude differences are very similar to those that were found in tCS by [9]. This indicates that the results of studying volume conduction effects in EEG source analysis also allow conclusions on the outcome of tCS simulations, and vice versa, and the theory for this relationship was presented here. Also, the AMG-CG FE solver method which was first introduced to EEG source analysis [26] can be efficiently used in tCS simulations. Lew and colleagues [25] demonstrated for FEM based EEG source analysis that, for a fixed accuracy, the AMG-CG solver achieved an order of magnitude higher computational speed than Jacobi-CG or IC(0)-CG, a result that can thus directly be transferred to the field of tCS simulation.

A last aim of this study was to compare tCS simulation in a geometry-adapted and a regular hexahedral FE approach in a highly realistic volume conductor model with white matter anisotropy. Significant changes in orientation up to 13.1 degrees and a decrease in current density amplitude of about 20 % occurred in the gray matter compartment when using the numerically more accurate geometry-adapted hexahedral approach as compared to the regular one. The effect sizes are similar to those of neglecting the distinction between skull compacta and spongiosa in skull modeling for tCS [9]. In clinical practice, the exact knowledge of current density orientation and magnitude is very important. Minor changes in the cortical current density vector field might even strongly influence the decision with respect to placement of the electrodes and/or strength of stimulation [4, 13]. Therefore, using a geometry-adapted FE approach might substantially increase accuracy and reliability of tCS simulation results and help to find optimized stimulation protocols. Current density amplitudes in the brain and CSF compartment were significantly reduced (up to 20 %) when using a geometry-adapted as compared to a regular hexahedral approach. Moreover, current density in the skin underneath the edges of the electrodes was decreased (up to 15%). Thus, commonly-used regular hexahedral FE approaches [10, 11, 7] might overestimate current densities in the brain and in the skin underneath the electrodes. In [9] we also demonstrated that strongest current densities always occur in the skin compartment underneath the edges of the electrodes, which might cause skin irritations or skin burn [56]. In summary, because skin and brain current magnitudes might have been overestimated in former regular hexahedral FE modeling studies, the proposed geometry-adapted approach suggests that higher stimulation magnitudes might be needed to accordingly modulate neural activity at brain level.

There are multiple limitations in our study that should be addressed. As often done in validation studies using multi-layer sphere models [15], the RDM and MAG error measures were only computed on the surface of the volume conductor model. However, these surface error measures for the electric potential distribution Φ were investigated for sources in the depth of the volume conductor and for the whole range of source eccentricities. Therefore, they reflect modeling deficits throughout the whole volume conductor and not only at the surface. For example, if the skull compartment had not been modeled accordingly, the RDM and MAG errors at the surface would clearly reflect this modeling deficit. Using reciprocity, the presented surface topography and magnitude error measures are thus indicators for the error in the electric potential at the source position for tCS modeling when using the same electrodes, too. However, these error measures are only reciprocal and not direct indicators. They are furthermore especially not direct measures for the error in the current density distribution ***J*** = *σ*∇Φ, for which we do not have analytical tools available. Nevertheless, the chosen error measures should still also be reasonable indicators for the errors in current density, as discussed in the following: In our chosen Lagrange-FEM approach, while the potential distribution Φ is continuous within the whole volume conductor, it is bending at the element boundaries, so that the resulting gradient of the potential (the electric field ∇Φ) might be discontinuous over element boundaries. At tissue boundaries with a large jump in the conductivity *σ* (e.g., from CSF to skull), this is also needed to keep the current as continuous as possible. The jump in ∇Φ thus mainly cancels the jump in tissue conductivities *σ* over this boundary to keep the current ***J*** = *σ*∇Φ mainly continuous. However, in a Lagrange FEM approach, the resulting current density ***J*** might still overall be discontinuous over the element boundaries. In contrast, in a Discontinuous Galerkin (DG) FEM approach [57], the current density distribution ***J*** is kept continuous within the whole volume conductor model, while the potential Φ is allowed to jump over element boundaries. In summary, Lagrange-FEM keeps the potential continuous, while the current might have discontinuities over element boundaries, and DG-FEM keeps the current continuous, while the potential might jump over element boundaries. In [54, 58], we implemented and compared both DG‐ and Lagrange-FEM approaches for tCS and the EEG forward problem and found significant differences between both approaches for both electric potential and current density only in case of models with thin skull compartments and insufficient resolution. These investigations thus show that our surface RDM and MAG error measures for the electric potential should be already reasonable indicators for the errors in current density, too, even if they are not direct measures and if they have to be seen in a reciprocal manner.

As a further limitation of our study, we clearly want to point out that in Figure 4, we only investigate the differences between the two hexahedral approaches. While the validations in the sphere models indicate that the accuracy of the geometry-adapted approach is better, we do not exactly know on which level of overall accuracy both the regular and the geometry-adapted hexahedral approaches are. For these reasons, computer simulations like in this study can only be a first step in validation, further validation studies in phantoms and/or animal models should thus be performed to compare simulated and measured current density distributions.

Finally, in our computer simulation study at hand, like in [59], the conductivity of the patches was modeled as saline, i.e., we also assumed well-soaked sponges in our simulations. However, the conductivity value of the electrode patches depends on the amount of saline in the patches. The contact impedance, surface of the stimulating electrodes and shunting currents influence our simulation results. In this paper, we modeled these aspects with the point electrode model (PEM) in combination with additional surface finite elements for the sponges, the so-called gap model [60]. They can, however, also be modeled using a complete electrode model (CEM) [61, 62, 63, 60]. In [60], it has been shown, that CEM and PEM only lead to small differences which are mainly situated locally around the electrodes and are very small in the brain region. Based on these results, the application of PEM and especially of the gap model like in the current study, being even closer to the CEM, is expected to result in negligible differences to the CEM and should thus provide a sufficiently accurate modeling of the current density within the brain region.

## Appendix

In the appendix, we will prove that Φ_*AA*_ ∈ ℝ^*S*–1^, i.e., the potential vector at the non-reference electrodes resulting from the adjoint approach, and *Φ*_*PI*_ ∈ ℝ^*S*–1^, the potential resulting from the partial integration approach in conjunction with the FE transfer matrix concept, are identical even for general head models.

### Lemma 1 (Matrix formulation for the adjoint approach solution vector).

Let (*b*_*i*_)_*i*=1, …, *S*–1_ = *B* ∈ ℝ^*N* × (*S*–1)^ be the matrix containing the right-hand sides of the adjoint approach. Let furthermore (*w*_*i*_)_*i*=1, …, *S*–1_ = *W* ∈ ℝ^*N*×(*S*–1)^ be the matrix containg the electric potential solution vectors of the adjoint approach. In addition, let us define *D*Ψ(*x*) := [∇*ψ*_1_(*x*), …, ∇*ψ*_*N*_(*x*)] ∈ ℝ^3×*N*^ with *ψ*_*i*_ being the FE ansatz functions and let us assume a current dipole at location *x*_0_ with dipole moment *q*. The potential Φ_*AA*_ is then given as

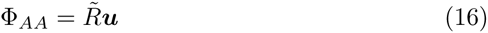

with

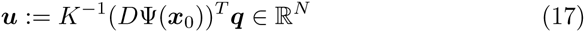

and *R*̃:= *B*^*T*^ ∈ ℝ^(*S*–1)×*N*^.

*Proof*. First, in equation (13), we can project the continuous potential function *w*_*i*_ : ℝ^3^ → ℝ in the finite element basis, i.e., 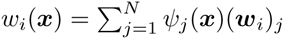. Therefore, the i^*th*^ entry of the adjoint approach potential vector Φ_*AA*_ ∈ ℝ^*S*–1^ for a single lead is given as

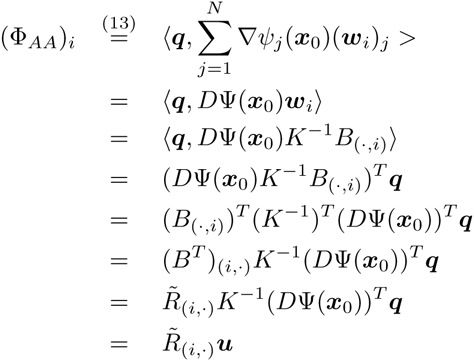

and thus Φ_*AA*_ = *R*̃*u*.

### Lemma 2 (Matrix formulation for the partial integration approach in conjunction with an FE transfer matrix).

Let *R* ∈ ℝ^(*S*–1)×*N*^ and be the restriction matrix from Equation (6). Then one obtains

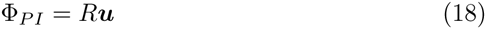

with the same *u* as in Lemma 1.

*Proof.* Φ_*PI*_ is simply given by

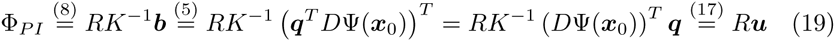

Theorem 1 relates the solution of the AA to the solution of the PI approach.

### Theorem 1.

Let Φ_*AA*_ ∈ ℝ^*S*–1^ and Φ_*PI*_ ∈ ℝ^*S*–1^ be the EEG forward potentials calculated with the adjoint approach and the partial integration approach in conjunction with the FE transfer matrix, respectively. Then, both EEG forward potential vectors are identical, whereas only the exact type of referencing is different.

*Proof.* Because *R*̂ *ε* ℝ^(*S*–1)×*N*^ and *R ε* ℝ^(*S*–1)×*N*^ only differ in column *i*_*ref*_ (the FE node of the reference electrode), we find

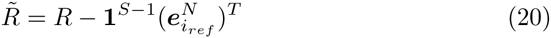

with 1^*S*–1^ *ε* ℝ^*S*–1^ a vector filled with 1 and 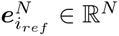 the unit vector with 1 only at position *i*_*ref*_. Therefore, the following equation holds:

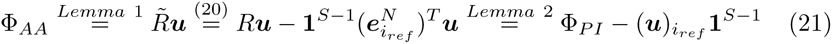

## Acknowledgement

This work has been supported by the priority program SPP1665 of the German Research Foundation, projects WO1425/5-1 (for SW, FL, JV, MB and CHW), HE3353/8-1 (for CSH) and EN533/13-1 (for GN).

## Footnotes

1 The SimBio-Vgrid mesh generator: www.rheinahrcampus.de/medsim/vgrid/index.html

2 SimBio: A generic environment for bio-numerical simulations, see www.mrt.unijena.de/simbio and www.simbio.de/.

3 www.fmrib.ox.ac.uk/fsl.

4 www.neuroscan.com/curry.cfm

5 SPM extension toolbox ACID (Algorithm HySCo): see www.fil.ion.ucl.ac.uk/spm/ext/ and www.diffusiontools.com/HySCo.html

6 Flexible Algorithms for Image Registration (FAIR): www.mic.uniluebeck.de/people/janmodersitzki/software/fair.html

